# MiR-30a-5p alters epidermal terminal differentiation during aging by regulating BNIP3L/NIX-dependent mitophagy

**DOI:** 10.1101/2021.10.25.465713

**Authors:** Fabien P Chevalier, Julie Rorteau, Sandra Ferraro, Lisa S Martin, Alejandro Gonzalez-Torres, Aurore Berthier, Naima El Kholti, Jérôme Lamartine

**Affiliations:** CNRS UMR 5305, Tissue Biology and Therapeutic Engineering Laboratory (LBTI), Lyon, France; Claude Bernard University Lyon 1, Villeurbanne, France; Gattefossé SA, 36 chemin de Genas - CS 70070 - 69804 Saint-Priest Cedex, France

**Keywords:** BNIP3L, miR-30a, mitophagy, keratinocyte, aging, mitochondria

## Abstract

Chronological aging is characterized by an alteration of the genes regulatory network. In human skin, epidermal keratinocytes fail to differentiate properly with aging, leading to the weakening of the epidermal function. MiR-30a is particularly overexpressed with epidermal aging, but the downstream molecular mechanisms are still uncovered. The aim of this study was to decipher the effects of miR-30a overexpression in the human epidermis, with a focus on keratinocyte differentiation. We formally identified the mitophagy receptor BNIP3L as a direct target of miR-30a. Using a 3D organotypic model of reconstructed human epidermis overexpressing miR-30a, we observed a strong reduction of BNIP3L expression in the granular layer. In human epidermal sections of skin biopsies from donors of different ages, we observed a similar pattern of BNIP3L decrease with aging. Moreover, human primary keratinocytes undergoing differentiation *in vitro* also showed a decreased expression of *BNIP3L* with age, together with a retention of mitochondria. Moreover, aging is associated with altered mitochondrial metabolism in primary keratinocytes, including decreased ATP-linked respiration. Thus, miR-30a is a negative regulator of programmed mitophagy during keratinocytes terminal differentiation, impairing epidermal homeostasis with aging.

## 1. Introduction

Aging is a natural and inexorable biological evolution associated with a progressive decline in tissue homeostasis. Skin is an excellent model of aging: during the natural aging process, the skin undergoes a typical age-related tissue dysfunction including epidermis atrophy, barrier dysfunction and delayed wound healing. This chronological and dynamic process starts in the mid-20s and is easily remarkable at the macroscopic level at advanced stage. However, early microscopic events such as changes in cell behaviour occur very early in the aging process and could be targeted to slow down the loss of tissue homeostasis. Although skin aging is strongly modulated by extrinsic factors, clues that a universal epigenetic program is driving the intrinsic tissue decline are converging in the literature. Indeed, multiple epigenetic changes are considered to be reliable hallmarks of tissue aging such as modification of DNA methylation patterns, histone posttranslational modifications as well as modulation of non-coding RNA expression (Pal and Tyler, 2016; Saul and Kosinsky, 2021). The latter is an emerging scientific domain in which an incredibly expanding number of studies have been published over the last decade. Indeed, the control of skin homeostasis depends largely on the fine-tuning of signalling pathways by long non-coding RNAs, circular non-coding RNAs and micro-RNAs (miRNAs) (Chevalier et al., 2020; Durante et al., 2021; Zeng et al., 2021).

We have previously conducted a genome-wide miRNA profiling in human primary keratinocytes from skin biopsies of young or elderly donors (Muther et al., 2017a). Thanks to this analysis we have identified miR-30a as the most over-expressed miRNA during the aging process in keratinocytes. The overexpression of miR-30 in young cells is sufficient to recapitulate in a 3D organotypic model the major functional defect of skin aging, namely the disrupted barrier function (Muther et al., 2017a). This functional alteration of the epidermis is likely due to increased apoptosis together with failure of the complete differentiation process. The epidermis is a multi-layered epithelium composed mainly of keratinocytes. Keratinocytes from the basal layer are proliferative cells that divide continuously to self-renew the pool of progenitor cells and sustain the daily need for committed cells. These committed keratinocytes will undergo progressive differentiation, in stages, along a vertical axis oriented outwards. At the terminal stage of differentiation, keratinocytes can be likened to mummified cells, called corneocytes, devoid of nucleus and organelles.

Macroautophagy (hereafter referred as autophagy) plays a crucial role for the terminal differentiation of keratinocytes, therefore its abnormal activity is associated with different cutaneous diseases (Monteleon et al., 2018). Autophagy is a dynamic process that degrades and recycles cellular components, such as misfolded proteins and damaged organelles. Multiple molecular complexes are involved in autophagy, but they will all lead to the sequestration of the targeted component within a double membrane autophagosome. This autophagosome will later fuse with a lysosome leading to a degradative autolysosome (Hill et al., 2021). In the epidermis, autophagy is constitutively active in the granular layer, where nuclei are cleared, leading to the terminal differentiation (Akinduro et al., 2016). In addition, mitophagy, a specialized type of autophagy that targets mitochondria for elimination, is a key step to activate the terminal differentiation of keratinocytes, through the BNIP3/BNIP3L (also known as NIX) pathway (Monteleon et al., 2018; Moriyama et al., 2014; Simpson et al., 2021). Interestingly, autophagy is a recurrent biological pathway targeted by miR-30 family in others systems, especially in many types of cancers (Pourhanifeh et al., 2020; Yang et al., 2021; Zhu et al., 2009). However, the functional role of miR-30a in skin and the consequences of its overexpression with aging are still uncovered in the literature. We hypothesized that miR-30a is a central regulator of mitophagy which will disturb the normal process of epidermal differentiation with aging. The present study was conducted on a 3D organotypic model of reconstructed epidermis and on human skin biopsies. We formally identified a new target of miR-30a and demonstrated that the mitochondrial dynamics is altered during differentiation along chronological aging.

## 2. Materials and Methods

### Cell culture

Human primary keratinocytes (HPK) were isolated in-house from skin biopsies as previously described (Muther et al., 2017b) or purchased from Lonza (Basel, Switzerland, #00192627). Skin biopsies were obtained from the DermoBioTec tissue bank at Lyon (Tissue Transfert Agreement n°214854) with the informed consent of adult donors or children’s parents undergoing surgical discard (non-pathological tissues from foreskin, ear, breast, or abdomen), in accordance with the ethical guidelines (French Bioethics law of 2004). HPK were cultured in KGM Gold medium (Lonza, #00192060) at 37°C and 5% CO_2_. The culture medium was renewed three times a week and cells were maintained no more than passage 4. The differentiation of HPK was induced by leaving the cells at confluency in KGM Gold medium. Human Embryonic Kidney (HEK293T) cells were cultured in Dulbecco’s modified Eagle’s Medium–Glutamax medium (DMEM) (Gibco, Waltham, MA, USA, #6196-026) 10% FBS at 37°C and 5% CO_2_ with renewal of culture medium three times a week.

### Transfection and luciferase assay

HPK at 70% confluency were transfected with miRNA mimics at 10nM: miR-30a-5p (Thermo Fisher Scientific, Waltham, MA, USA, #4464066, assay ID: MC11062), miR-30a-3p (Thermo Fisher Scientific, #4464066, assay ID: MC10611) and miRNA negative control (Thermo Fisher, #4464058), using RNAiMax (Thermo Fisher Scientific, #13778075) according to manufacturer’s instructions. The cells were processed for RT-qPCR analysis of *BNIP3L* expression 24h after transient transfection. Partial sequence of *BNIP3L* 3’UTR was cloned downstream of a firefly luciferase reporter plasmid (VectorBuilder, Neu-Isenburg, Germany, #VB191204-1713qzf). The plasmid was then submitted to site-directed mutagenesis using the In-Fusion technology (Takara Bio Europe, Saint-Germain-en-Laye, France, #638909) with inverse PCR (see supplementary Table 1) according to manufacturer’s instructions. The characterization of the resultant plasmids was performed by Sanger sequencing (Biofidal, Vaulx-en-Velin, France). An additional renilla luciferase reporter plasmid was generated to serve as an internal standardizer (VectorBuilder, # VB191205-1069fzu). HEK293T cells at 70% confluency were co-transfected with the firefly luciferase reporter plasmids at 400 ng/mL, the renilla luciferase reporter at 0,4 ng/mL and miRNA mimics at 10 nM using TransIT-X2 (Mirus Bio, Madison, WI, USA, #MIR 6000) according to manufacturer’s instructions. 18h after transfection, the luciferase activity was measured using the renilla-firefly luciferase dual assay kit (Thermo Fisher, #16186) in a microplate reader (Infinite M1000, Tecan, Männedorf, Switzerland).

### Reconstructed human epidermis (RHE) production

RHE were prepared as described previously in Muther et al., 2017 [1]. Briefly, 3.10^4^ primary fibroblasts were seeded on the outer face of polycarbonate membrane of cell culture inserts (Millipore, Sigma-Aldrich, Saint-Louis, MO, USA) and cultured for two days. Then 3.10^5^ primary keratinocytes, infected by pSLIK Venus control or pSLIK Venus miR-30a, were seeded on the inner face of inserts. Three days after, differentiation and stratification were induced when keratinocytes were placed at air-liquid interface. The culture was maintained for 11 days and culture medium was changed every day during the immersion phase. Doxycycline (Sigma-Aldrich) at 0.1 μg/mL was added for the time of the protocol to induce miR-30 over-expression.

### Immunofluorescence

Human paraffin embedded tissue microarray (TMA: #SK2444A and #SKN1001) were purchased from US Biomax (Derwood, MD, USA). Antigen retrieval (10mM sodium citrate buffer, 0.05% Tween20) was performed at 95°C during 20 min before immunostaining. RHE were prepared for immunofluorescence as previously described (Muther et al., 2017b). The following steps are similar for both TMA and RHE sections. After dewaxing and rehydration, tissue sections were permeabilized using PBS 0.1% Triton X-100, 0.1M glycine during 10 min at room temperature (RT). Samples were then blocked with PBS containing 5% goat serum, 2% BSA and 0.1% Tween20 for at least 1h at RT. After washing steps, primary antibodies were incubated overnight at 4°C (supplementary Table 2). After washings, secondary antibodies were incubated 45min at RT and nuclear staining was performed using ProLong^™^ Glass Antifade Mountant with NucBlue^™^ (Thermo Fischer Scientific). Negative controls were performed by omitting primary antibodies. Image were visualized using High Content Screening Yokogawa CQ1 microscope (Yokogawa, Tokyo, Japan), digitalized using sCMOS camera (Olympus, Hamburg, Germany) and analysed using ImageJ software (version 2.1.0/1.53c).

### Total RNA/DNA isolation and real-time quantitative PCR

Total RNA and DNA were isolated using Quick-DNA/RNA^™^ Miniprep kit (Zymo research, Mülhauser, Germany) according to manufacturer’s instruction. First, mRNA were reverse-transcribed into cDNA using PrimeScript^™^ RT reagent kit (Takara, Shiga, Japan) and analysed on real-time qPCR using SYBR^®^ Premix ExTaqII (Takara) on an AriaMx Realtime PCR system (Agilent Genomics, Santa Clara, CA, USA). Results were normalized to *TBP* and *RPL13A* housekeeping gene expression levels, using the 2^-ΔΔ^Ct quantification method. Secondly, relative mitochondrial DNA content was calculated as the mean ratio of two mitochondrial genes copy number (*ND1* and *TL1*) to single-copy nuclear genes (*HBB* and *SERPINA1*) using the 2^-Δ^Ct quantification method. All primers are listed in the supplementary Table S1.

### Seahorse analysis

Oxygen consumption rate (OCR) was measured with a Seahorse XF extracellular flux analyzer according to the manufacturer’s instructions (Agilent). Primary keratinocytes were seeded at 60,000 cells per well (4 wells as technical replicate/cell type) in a 24-well cell culture microplate and incubated overnight at 37°C in 5% CO_2_. Culture medium was replaced with XF assay medium supplemented with 10mM glucose (XF Glucose solution, 103577-100, Agilent), 1 mM pyruvate (XF Pyruvate solution, 103578-100, Agilent) and 2 mM glutamine (XF Glutamine solution, 103579-100, Agilent) and cells were incubated in the absence of CO_2_ for 45 min before measurement. OCR was determined before injection of specific metabolic inhibitors and after successively adding 1.5 μM oligomycin, 1 μM FCCP, and 0.5μM rotenone/antimycin A (Sigma-Aldrich). Wave software was used to analyse seahorse measurements.

### Statistical analysis

Data are expressed as mean or median ± SD. Statistical significance was calculated by Student’s t-test, one-way analysis of variance (ANOVA), two-way analysis of variance (ANOVA2), or Pearson correlation using Prism software (version 7.0, GraphPad Software, San Diego, CA, USA). Mean differences were considered statistically significant when p < 0.05. * p < 0.05, ** p < 0.01, *** p < 0.001, **** p < 0.0001.

## 3. Results

### 3.1. BNIP3L is a new identified target of miR-30a-5p

Because miR-30a is already well known to regulate autophagy through targeting *BECLIN1* or *ATG5* mostly in cancer cells (Pourhanifeh et al., 2020; Yang et al., 2021; Zhu et al., 2009), we chose to focus on putative targets involved more specifically in mitophagy. Using TargetScan (http://www.targetscan.org/vert_72/), we found that miR-30a-5p has two highly conserved putative binding sites in *BNIP3L* 3’UTR at positions 2350-2357 and 2651-2657 (Figure 1A). In addition, another putative binding site of miR-30a-5p (position 1126-1132) and two putative binding sites of miR-30a-3p (positions 1082-1088 and 2059-2065) were also found, even though the conservation of these sites among vertebrates is less important. Since the both strands of miR-30a are overexpressed in keratinocytes from aged skin, we tested the *in vitro* ability of miRNA mimics to decrease the mRNA levels of *BNIP3L* in keratinocytes (Figure 1B)., Both mimics of miR-30a-3p and miR-30a-5p induced a significant decrease of *BNIP3L* mRNA levels by 28% and 36% respectively, 24h after transient transfection of proliferating human keratinocytes.

**Figure 1.**
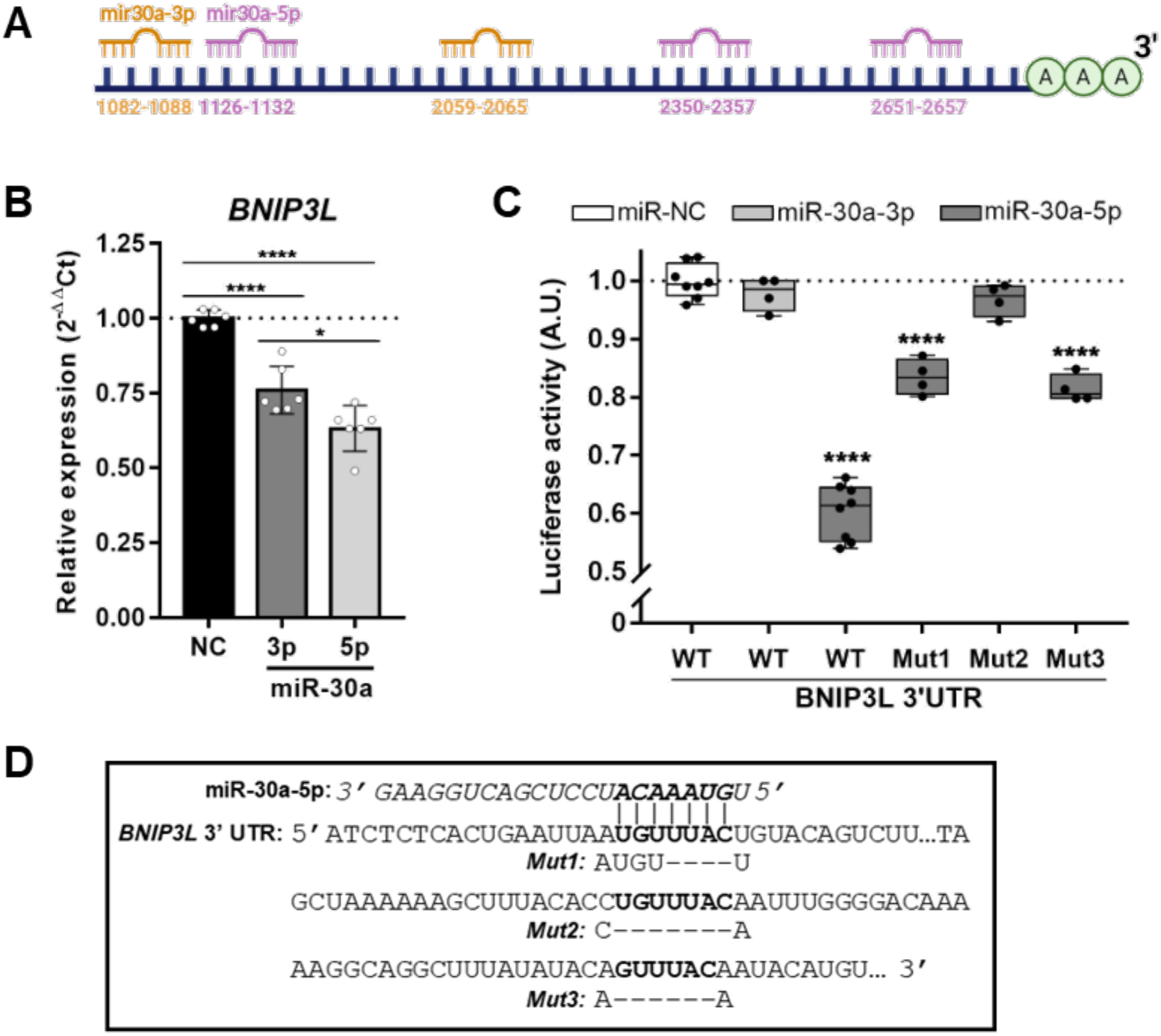
miR-30a interactions with *BNIP3L*. (**A**) Schematic of the *BNIP3L* 3’UTR region showing the putative miR-30a binding sites; (**B**) Relative mRNA expression of *BNIP3L* in keratinocytes 24h after transient transfection of miR-30a-3p or miR-30a-5p mimics or a scrambled sequence (NC: Negative Control). qPCR analysis was normalized to *TBP* and *RPL13A* housekeeping genes using the 2^-ΔΔ^Ct quantification method (mean ± SD, n=6; *p<0.05 and ****p<0.0001). Exact p-values were determined using the One-way ANOVA and Tukey post-hoc tests. (**C**) Normalized luciferase activity (red firefly/green renilla) after co-transfections of HEK293T cells with the Wild-Type (WT) or Mutant (Mut) reporter plasmids and miRNA mimics (median ± SD, n=4 or n=8; ****p<0.0001). Exact p-values were determined using the One-way ANOVA and Tukey post-hoc tests. (**D**) Schematic of a portion of *BNIP3L* 3’UTR region with the three putative complementary pairing sites with miR-30a-5p. The mutated sites, obtained by inverse PCR and characterized by Sanger sequencing, are also indicated below the original sequence.

To further confirm the direct targeting of *BNIP3L* by the miR-30a, a portion of its 3’UTR sequence containing the five putative binding sites was cloned into a luciferase reporter plasmid. In combination with transient transfection of either miR-30a-3p or miR-30a-5p mimics in HEK293T cells, we observed a strong reduction (−40%) of the luminescence signal only in presence of the miR-30a-5p strand (Figure 1C). Thus, it appears that only miR-30a-5p is directly targeting the *BNIP3L* 3’UTR and that the effect of miR-30a-3p previously observed in primary keratinocytes is likely related to a secondary event. We then selectively mutated the three putative binding sites of miR-30a-5p individually. Sanger sequencing showed a partial deletion of four nucleotides out of seven of the first binding site (1126-1132) and complete deletion of all seven nucleotides from the two others binding sites (2350-2357 and 2651-2657) (Figure 1D). Mutants 1 and 3 partially rescued the luciferase activity (from −40% to −17% and from - 40% to −19%, respectively), suggesting that these two sites only account for a partial activity of miR-30a-5p on *BNIP3L* 3’UTR. Surprisingly, the luciferase activity was almost fully restored (from −40% to −3%) when the intermediate binding site was mutated, suggesting that this single site is sufficient for miR-30a-5p activity. These apparent conflicting data may reflect of non-canonical miRNA-target interactions, independently of the seed complementary sequence (Seok et al., 2016), in the proximity of the both sites 1126-1132 and 2651-2657.

### 3.2. miR-30a-5p abolishes BNIP3L expression in the granular layer of RHE

To confirm the regulation of *BNIP3L* by miR-30a in a more relevant system, we took advantage of an organotypic model of RHE generated from HPK overexpressing miR-30a. Using three different human donors, we showed that keratinocytes normally express BNI3PL in the granular layer of the epidermis (Figure 2A), in accordance with its described function in the degradation of mitochondria during the cornification (Simpson et al., 2021). Strikingly, the overexpression of miR-30a in keratinocytes induced a strong reduction of BNIP3L staining in the granular layer. The reminiscence of BNIP3L protein is likely due to cell heterogeneity in overexpression of miR-30a, since we used primary human keratinocytes that did not undergo through a selection pressure. Nevertheless, we measured the percentage of the granular layer length with positive staining for BNIP3L with or without miR-30a over-expression. Under normal conditions, 88% of the granular layer is positive for BNIP3L, whereas in the presence of high levels of miR-30a only 18.5% is stained for this mitophagy protein (Figure 2B).

**Figure 2.**
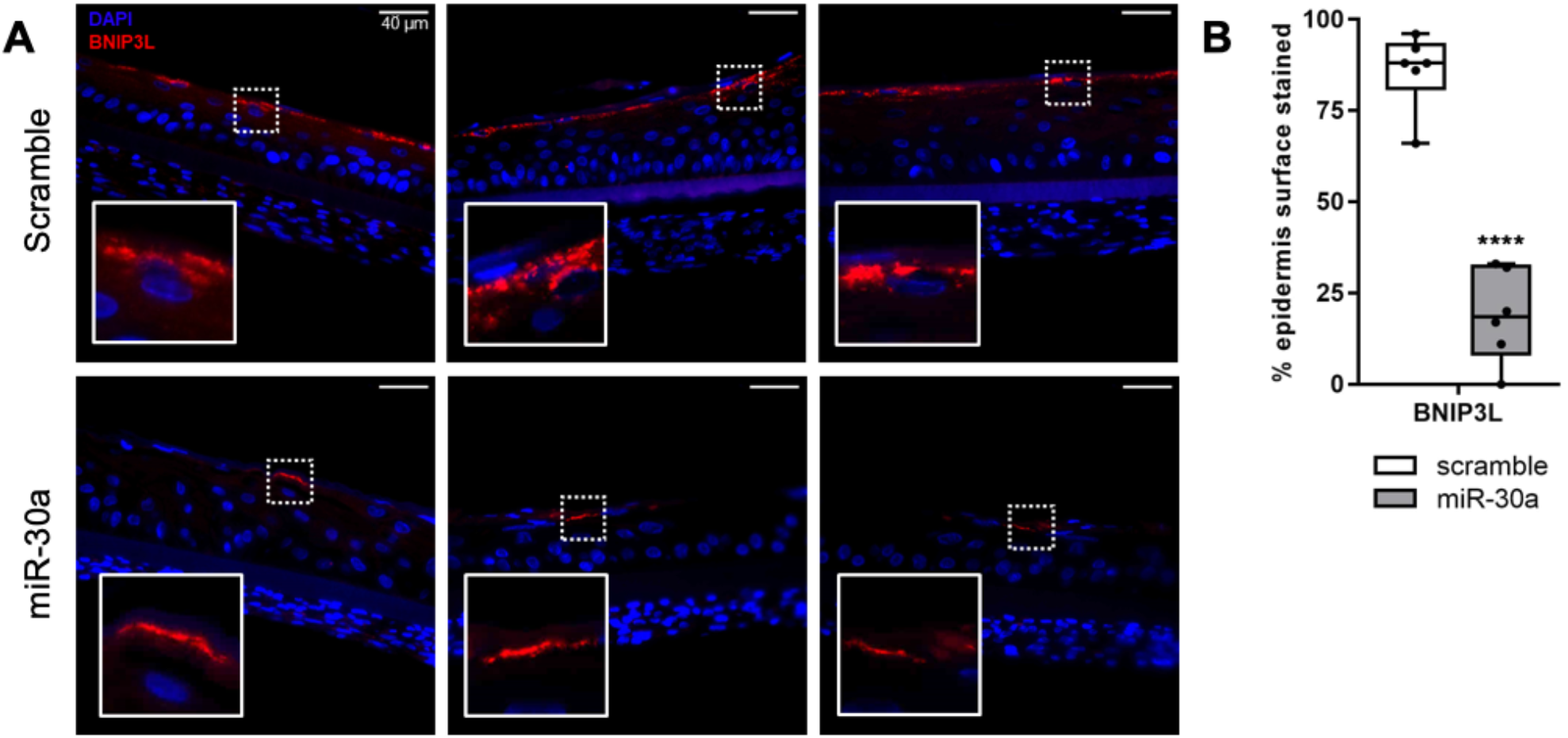
Effect of miR-30a on BNIP3L expression in the granular layer of reconstructed epidermis. (**A**) Immunofluorescent staining of BNIP3L in reconstructed human epidermis overexpressing or not miR-30a. Representative photographs of 3 independent replicates are shown. (**B**) Quantification of the surface of the epidermis with a positive signal for BNIP3L in each condition (median ± SD, n=3; ****p<0.0001). Exact p-value was determined using the Student’s *t*-test.

### 3.3. BNIP3L expression decreases with chronological aging in human epidermis

Since miR-30a expression is known to increase with chronological aging in the epidermis, we used a TMA to screen the expression of its target BNIP3L in human skin biopsies of different ages. We defined two groups based on the age of the donor and on the structure of the skin: adult individuals (n=11, from 19 to 42 years old, mean age at 34,4) vs aged individuals (n=10, from 61 to 78 years old, mean age at 68,3). Indeed, skin aging is characterized in terms of tissue morphology by epidermal atrophy and flattening of the dermo-epidermal junction. Illustrative examples of skin biopsies from three adults (Figure 3A, top panel: 38, 35 and 42 years old) and three aged individuals (Figure 3A, bottom panel: 61, 71 and 78 years old) show the pattern of BNIP3L staining together with KRT14, a specific marker of basal undifferentiated keratinocytes. We found that BNIP3L is highly expressed in the upper layers of epidermis of the adult’s group. In addition, BNIP3L staining is often delimitating cornified cells, that are devoid of a nucleus (Figure 3A, inserts). With aging, we observed a strong decline of the BNIP3L signal along with a substantial decline in the number of enucleated keratinocytes. We evaluated the correlation between the percentage of positive cells for BNIP3L staining and age and the correlation of the normalized signal intensity per epidermal area with age (Figure 3B and 3C). Using these two correlations, we found equivalent Pearson coefficients with r = −0.6542 (R^2^ = 0.428, p=0.0024) and r = −0.6198 (R^2^ = 0.3841, p=0.0036) respectively. These mathematical simulations strongly suggest that the chronological aging negatively affects the expression of BNIP3L in human epidermis.

**Figure 3.**
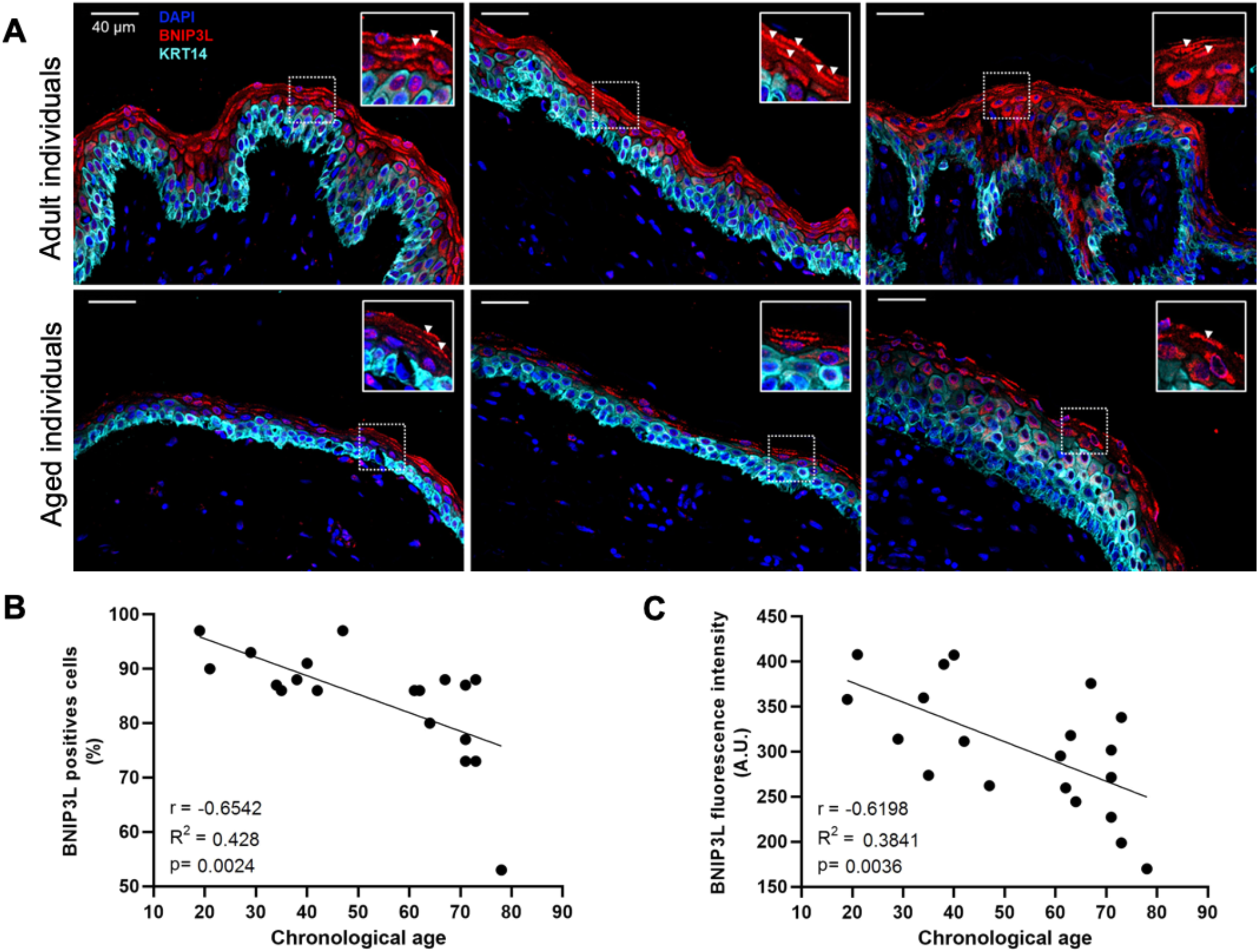
BNIP3L expression in human skin biopsies from individuals at different ages. (**A**) Immunofluorescent staining of BNIP3L and KRT14 in human skin sections. Representative photographs of 6 indivuals from two different age groups: adult, or aged people (top, from left to right: 38, 35 and 42 years old; bottom, from left to right: 61, 71 and 78 years old). Inserts are a magnification of the selected area showing BNIP3L-positive cells (white arrowheads) (**B**) Mathematical correlation between the percentage of BNIP3L-positive cells from the whole epidermis and the chronological age of the individual from which the skin biopsy has been sampled (n=21). Exact p-value was determined using the Pearson correlation test. (**C**) Mathematical correlation between the total BNIP3L fluorescence intensity from the whole epidermis and the chronological age of the individual from which the skin biopsy has been sampled (n=21). Exact p-value was determined using the Pearson correlation test.

### 3.4. Aging is associated with alterations of keratinocyte terminal differentiation, BNIP3L expression and mitochondrial elimination

In order to confirm that BNIP3L is differently regulated with aging during epidermis differentiation, we used an *in vitro* 2D model of differentiation with primary keratinocytes isolated from three cohorts of age: young individuals (n=6, from 3 to 10 years old, mean age at 5), adult individuals (n=6, from 26 to 46 years old, mean age at 36,5) and aged individuals (n=5, from 68 to 92 years old, mean age at 79). Considering the young cohort as a reference, we examined RNA expression levels of several markers along a nine-day differentiation kinetic (Figure 4A).

**Figure 4.**
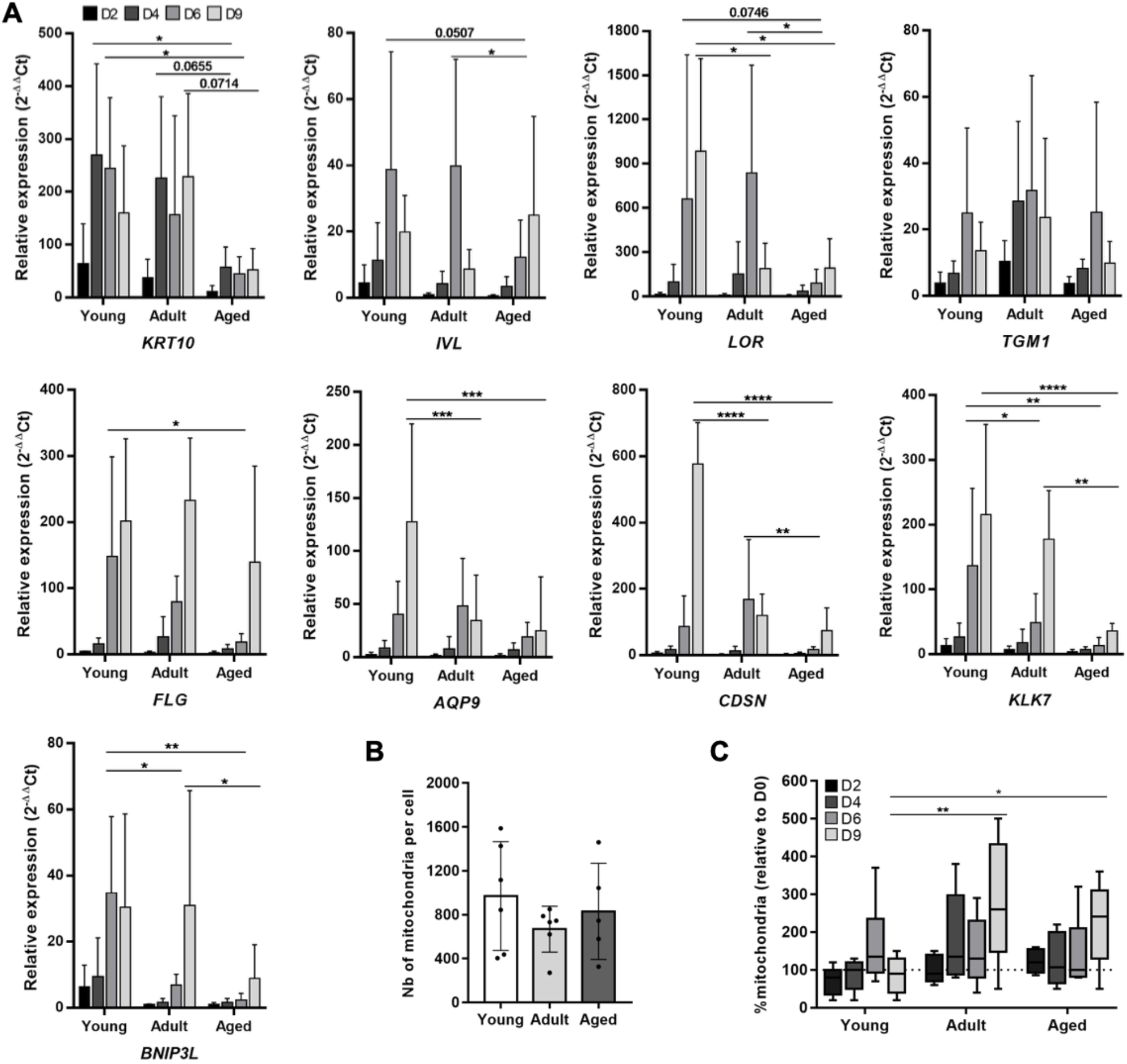
Human keratinocyte differentiation, *BNIP3L* expression and mitochondria retention *in vitro* with aging. (**A**) Relative mRNA expression of differentiation markers and *BNIP3L* in keratinocytes during a kinetic of differentiation in 2D culture. Three groups of different age were tested (young group: n=6, from 3 to 10 years old, mean age at 5; adult group: n=6, from 26 to 46 years old, mean age at 36,5; aged group: n=5, from 68 to 92 years old, mean age at 79) and followed over the time (0, 2, 4, 6 and 9 days after reaching confluency). qPCR analysis was normalized to *TBP* and *RPL13A* housekeeping genes (mean ± SD; \**p*<0.05, \*\**p*<0.01, \*\*\**p*<0.001 and \*\*\*\**p*<0.0001). Exact p-values were determined using the Two-way ANOVA and Tukey post-hoc tests. (**B**) Content of mitochondria per cell estimated by qPCR at day 0. qPCR analysis was performed by normalizing the relative content of mitochondrial genes (*ND1* and *TL1*) with the content of nuclear genes (*HBB* and *SERPINA1*) using the 2^-Δ^Ct quantification method in the three groups (mean ± SD; young group: n=6; adult group: n=6; aged group: n=5). (**C**) Evolution of the content of mitochondria expressed as a percentage of the quantity at day 0 (median ± SD; \**p*<0.05, \*\**p*<0.01). Exact p-values were determined using the Two-way ANOVA and Tukey post-hoc tests.

The commitment of keratinocytes to differentiation, illustrated by the expression of the early marker *KRT10*, was not found to be affected between young and adult groups, whereas keratinocytes from the aged group reduced *KRT10* expression to an approximately 5-fold extent. With the progression of the differentiation, we noticed that the expression levels of most markers (*IVL*, *LOR*, *TGM1*, *FLG*) in the adult group were not quite different from those in the young group. Exception is made for the expression of *LOR* which dropped at day 9 in the adult group while it was maintained in the young group. Considering the keratinocytes in the aged group, we observed both a delay in the intensity of expression and a strong decrease of the maximal level of expression (*IVL*, *LOR*, *FLG*), except for *TGM1* which is homogeneously expressed within the three age groups. However, the expression of the terminal differentiation genes, *KLK7*, *AQP9* and *CDSN*, were significantly decreased in both the adult group and the aged group, as compared to the young one. Interestingly, we found that *BNIP3L* is normally overexpressed from day 6 of differentiation, a time corresponding to the switch from late differentiation to terminal differentiation, as illustrated by the concurrent increase of *KLK7*, *AQP9* and *CDSN*. The pattern of *BNIP3L* expression during differentiation in keratinocytes somehow summarized the profile of the different age groups. The maximal expression level was reached from day 6 and maintained at day 9 in keratinocytes from young donors. In cells from adult donors, we observed a delayed maximal expression level, at day 9, but with a similar intensity of expression than the young group. Finally, keratinocytes from aged donors also expressed *BNIP3L* at the maximal at day 9, but with a significant reduction in the intensity compared with young and adult groups. This result is in accordance with the variation of BNIP3L immunostaining during aging in the human epidermis sections. We summarized the moment and the intensity of the maximal expression levels (relative to D0) for each gene in each group in Table 1.

**Table 1.** Moment and intensity of the maximal expression levels of differentiation markers in human keratinocytes (relative to D0).

|  | Gene | *KRT10* | *IVL* | *LOR* | *TGM1* | *FLG* | *AQP9* | *CDSN* | *KLK7* | *BNIP3L* |
| --- | --- | --- | --- | --- | --- | --- | --- | --- | --- | --- |
| Groups of age | Young | D6 | D6 | D9 | D6 | D9 | D9 | D9 | D9 | D6 |
|  |  | 244.7 | 38.7 | 984 | 24.9 | 201.4 | 127.7 | 576.8 | 215.8 | 34.7 |
|  | Adult | D9 | D6 | D6 | D6 | D9 | D6 | D6 | D9 | D9 |
|  |  | 228.5 | 39.8 | 836 | 31.8 | 232.8 | 48.2 | 167.7 | 177.7 | 30.9 |
|  | Aged | D9 | D9 | D9 | D6 | D9 | D9 | D9 | D9 | D9 |
|  |  | 52.4 | 25 | 189.5 | 25.2 | 139.7 | 25.2 | 74.1 | 35.8 | 8.9 |

Considering the role of BNIP3L in mitochondrial fragmentation associated to epidermis terminal differentiation (Simpson et al., 2021), we then evaluated the relative amount of mitochondria in the three cohorts of age during the 2D differentiation kinetic. Remarkably, we observed a strong inter-individual variation in the number of mitochondria per proliferating keratinocytes (D0), with no difference in the overall median between the groups of age (Figure 4B). In the young group, the number of mitochondria increased progressively to reach a peak at day 6, and then dropped at day 9 (Figure 4C; relative to D0: D2=80%, D4=100%, D6=135%, D9=90%). We believe that the increase of mitochondria is necessary during differentiation to keep up with high metabolic demand before the terminal differentiation. In 2D, there was no complete elimination of the organelles and nucleus because the final cornification step requires to be in contact with air. In the adult group, we observed an early increase of the mitochondria content, as soon as day 4, which is maintained at day 6 and even pursued at day 9 (relative to D0: D2=110%, D4=135%, D6=125%, D9=260%). The evolution of mitochondria in keratinocytes from the aged group followed the same profile as those of the adult group, with a milder intensity but with a retention of mitochondria at day 9 as well (relative to D0: D2=120%, D4=107%, D6=100%, D9=241%). These results overlap with the alteration of *BNIP3L* expression from day 6 in the adult and aged groups and may reflect a defective mitophagy during terminal differentiation of keratinocyte. However, the increased expression of *BNIP3L* at day 9 in the adult group suggests that either the induction of *BNIP3L* was too late for proper mitochondrial elimination, or that additional mechanisms of mitochondrial dynamics are also involved.

### 3.5. Aged keratinocytes display mitochondrial metabolic defects

As mitophagy serves as a quality control process of functional mitochondria, compromised mitophagy may alter the mitochondrial metabolism. Therefore, we measured the OCR in primary keratinocytes from the three groups of age. We found that the basal OCR was significantly decreased in the adult group (−22.7%) and the aged group (−32.1%), as compared to the young one (Figure 5A). Using the ATP-synthase inhibitor oligomycin, we found that ATP-linked respiration was also significantly decreased by 23.5% in the adult group and 30.6% in the aged group (Figure 5B). Next, we added FCCP, a protonophore which collapses the proton gradient across the mitochondrial inner membrane. This drug forces the electron transport chain to function at its maximal rate. Even if the maximal respiration capacity was identically altered than the basal respiration rate in the adult or the aged group (Figure 5C), the reserve capacity, expressed as the percentage of the basal rate, was identical within the three groups (Figure 5D). Lastly, to completely shut down the electron transport chain function, we added antimycin A and rotenone, two inhibitors of complex III and I respectively. Consequently, the remaining OCR is associated to non-mitochondrial respiration and allows the determination of the proton leak during mitochondrial respiration. The non-mitochondrial oxygen consumption, related to the activity of diverse desaturase and detoxification enzymes, seemed altered with aging as well. In the adult group, we observed a 30% decrease, albeit not significant (p=0.0995), whereas in the aged group this non-mitochondrial use of oxygen was decreased by half (Figure 5E). Finally, as for the others parameters, the proton leak is decreased with aging, but it invariably accounts for about 25% of the basal respiration in all groups (Figure 5F). This result suggests that the proton permeability of the inner mitochondrial membrane is not affected with age. Contrariwise, and considering that the number of mitochondria did not vary with age (Figure 4B), it seems that the mitochondrial metabolic activity is decreased with aging, which likely results in a decreased ATP synthesis.

**Figure 5.**
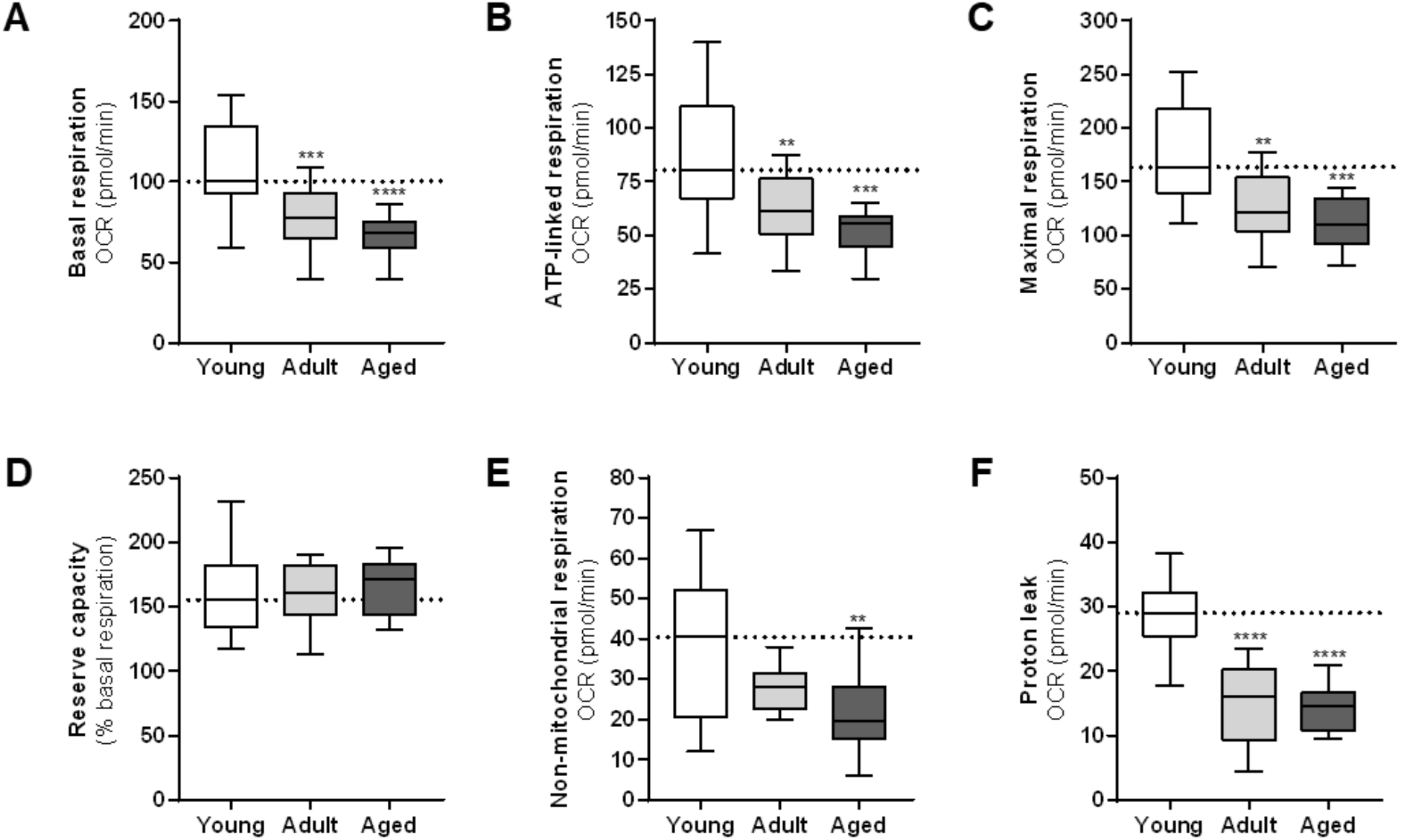
Mitochondrial metabolic activity of keratinocytes from different age groups. Major aspects of mitochondrial coupling and respiratory control were measured using the Seahorse Bioanalyzer and determined by the sequential additions of oligomycin, an ATP synthase inhibitor, FCCP, a protonophoric uncoupler, and rotenone and antimycin A, two inhibitors of the electron transport chain. Basal respiration (A), ATP-linked respiration (B), maximal respiration (C), reserve capacity (D), non-mitochondrial respiration (E) and proton leak (F) were determined by measuring the Oxygen Consumption Rate (OCR) in the culture media (median ± SD; young group: n=6; adult group: n=6; aged group: n=5; \*\**p*<0.01, \*\*\**p*<0.001 and \*\*\*\**p*<0.0001). Exact p-values were determined using the One-way ANOVA and Tukey post-hoc tests.

## 4. Discussion

BNIP3L (BCL2 Interacting Protein 3 Like)/NIX is a mitochondrial outer membrane protein of the BH3-only member of the BCL2 family, initially identified as a pro-apoptotic inducer (Chinnadurai et al., 2008; Ney, 2015). In addition to its role in the regulation of mitochondrial-dependent apoptosis, BNIP3L has also been identified as a critical autophagy receptor for the selective clearance of mitochondria in mammalian cells. For instance, BNIP3L is essential for mitochondria removal during terminal differentiation of multiple cell types including erythroid cells (Sandoval et al., 2008; Schweers et al., 2007), ocular lens fiber cells (Brennan et al., 2018), cardiac progenitor cells (Lampert et al., 2019) and optic nerve oligodendrocytes (Yazdankhah et al., 2021). A recent report showed that BNIP3L is also critical for the terminal differentiation of epidermal keratinocyte (Simpson et al., 2021). Indeed, the authors used CRIPSR-Cas9 to knock-out BNIP3L in immortalized human keratinocytes (hTERT cell line) and generated reconstructed epidermis from these modified cells. They obtained a stratified epithelium but noticed maturation defects. In details, they described that BNIP3L-deficient keratinocytes failed to undergo through complete cornification, with retention of mitochondria in the uppermost layers. In the present study, we found that BNIP3L is a direct target of miR-30a and we have previously reported that miR-30a expression levels substantially increase with aging in human keratinocytes (Muther et al., 2017b). Skin aging is associated to epidermal differentiation defects (Rinnerthaler et al., 2013; Wang et al., 2020), and we previously correlated that overexpression of miR-30a alters the differentiation process of keratinocytes. Indeed, using an organotypic model of reconstructed human epidermis (RHE), we have demonstrated that miR-30a overexpression strongly downregulates the differentiation markers such as KRT1, KRT10, IVL and LOR, and decrease the efficacy of the barrier function (Muther et al., 2017b). Accordingly, we found here that BNIP3L expression is strongly reduced with aging in human skin. Using a 2D model of keratinocyte differentiation, we also observed a retention of mitochondria with aging, thus imitating the findings of the study with BNIP3L-deficient keratinocytes (Simpson et al., 2021). Besides the reduction of BNIP3L expression with age and its role in the differentiation process, we also found here that the mitochondrial metabolism is affected during chronological aging.

Mitochondrial metabolism is the main source of cellular energy by generating high levels of ATP. Besides the role of ATP in providing energy for many cellular processes, it has been demonstrated that inhibition of ATP synthase blocks the differentiation of keratinocytes (Xiaoyun et al., 2017). In this study, exposure of the HaCaT cell line to oligomycin induced a significant decrease of the intracellular ATP content while extracellular ATP increased in parallel. Whereas no effect on keratinocytes proliferation was noted, the decreased intracellular ATP concentration was associated with an inhibitory effect on involucrin expression. Using a confluence-dependent differentiation model, as we used in the present study, the authors also showed that the ATP5B subunit of ATP-synthase was induced at days 6 and 8 of differentiation, both at the mRNA and protein levels. Even if the molecular mechanisms are not elucidated yet, these results strongly suggest that the intracellular ATP produced by mitochondrial metabolism is closely related to the terminal differentiation of keratinocytes. Accordingly, we observed a significant reduction of the oxygen consumption associated to ATP synthesis in keratinocytes from aged donors, together with alteration of their terminal differentiation.

Interestingly, a recent study has pointed out that mitophagy plays a key role in the regulation of bioenergetics in neurons. Using a mouse model of Alzheimer’s disease (AD), they observed a deficiency in energetic enhancement upon oxidative phosphorylation stimulation in brains (Han et al., 2021). In their model, the high energetic demand normally induced mitochondrial turnover by mitophagy, which is impaired in AD. Since the onset of AD is strongly associated with aging, we can hypothesize that the observed deficit in oxidative phosphorylation in keratinocytes is also linked to defective mitophagy with aging. In the skin, another study has found that ATP content was higher in primary dermal fibroblasts from centenarians, as compared to fibroblasts isolated from adult (about 27 years old) and aged (about 75 years old) human subjects, even though centenarians also presented oxidative phosphorylation defects (Sgarbi et al., 2014). The preservation of an adequate ATP production was associated with an increased mitochondrial mass together with re-arrangement of the mitochondrial network. Mitochondrial dynamics, a balance between mitochondrial fission and fusion, is critical to sustain functional mitochondria. Fission allows to create new mitochondria, but it also contributes to quality control by removing the damaged components. Conversely, fusion relieves stress by mixing the contents of damaged and undamaged mitochondria. In dermal fibroblasts from centenarians, the authors found more hyperfused and elongated mitochondria compared to younger subjects. They concluded that mitochondrial fission was reduced in these individuals undergoing what we might call “successful aging”. Since our study did not include cells from centenarians, it would be interesting to verify whether the remodelling of the mitochondrial network observed in dermal fibroblasts also occurs in keratinocytes from long living individuals.

Finally, even if changes in mitochondrial dynamics have been shown to contribute to other age-related disorders, such as cardiac disorders, neurodegenerative diseases and sarcopenia (reviewed in (Srivastava, 2017)), mitophagy and the balance between fission and fusion, are not the only processes to achieve mitochondrial quality control. In response to oxidative stress, cells initiate an external release of mitochondria (Liu et al., 2020). This mechanism occurs as an early response and precedes mitophagy in HeLa cells. To insure cellular homeostasis, damaged mitochondria or their constituents will be transferred out of the cells in free form or encapsulated in extracellular vesicles (EVs). This second process functions in complementarity with mitophagy: it is repressed when mitophagy is efficient, whereas it is activated to compensate lack of mitophagy (Choong et al., 2020). In the present study, we demonstrated that aged cells, which highly express miR-30a, exhibit a defective BNIP3L-dependent mitophagy in the last steps of keratinocyte differentiation. In an aging context, the external mitochondrial release by keratinocytes does not seem to be enough to compensate defective mitophagy and restore mitochondrial homeostasis. This statement is in good agreement with a previous study which reported that the amount of mitochondrial DNA released in EVs decreases with age (Lazo et al., 2021). Thus, mitochondrial dysfunction observed in skin aging may be due to breakdown in the two mitochondrial quality control systems. Moreover, EVs, as key vectors of intercellular communication (Yáñez-Mó et al., 2015), allows the transfer of intact mitochondria or functional mitochondrial proteins and DNA to other cells. Several studies have analysed the effect of EVs containing mitochondrial components on other cells (Noren Hooten and Evans, 2021) and it has notably been shown that EVs from older individuals affect mitochondrial function of HeLa cells with a significant decrease of basal respiration (Lazo et al., 2021). Interestingly, we found in this study that keratinocytes from adults have an intermediate profile. They had higher expression levels of BNIP3L than keratinocytes from aged individuals, but they already displayed the equivalent altered mitochondrial metabolic activity, including lower basal respiration, as compared to cells from young individuals. Thus, one can hypothesize that extracellular release of mitochondria plays an important role in the propagation of aging signals in skin tissue, leading from an adult phenotype to an aged one. Additional studies are needed to further validate this assumption.

## Supplementary Materials

Table S1: List of primers used for qPCR and PCR assays, Table S2: List of antibodies used for immunofluorescence.

## Author Contributions

Conceptualization, F.P.C., J.R. and J.L.; methodology, F.P.C., J.R., A.G-T., A.B., N.E.; validation, F.P.C., J.R., S.F. and L.S.M.; formal analysis, F.P.C., J.R. and L.S.M.; investigation, F.P.C., J.R., S.F., L.S.M., A.G-T., A.B. and N.E.; resources, F.P.C. and J.L.; writing–original draft preparation, F.P.C., J.R. and J.L.; visualization, F.P.C., J.R., L.S.M. and J.L.; supervision, F.P.C. and J.L.; project administration, F.P.C.; funding acquisition, F.P.C. and J.L. All authors have read and agreed to the published version of the manuscript.

## Acknowledgments

We acknowledge the contribution of SFR Biosciences (UAR3444/CNRS, US8/Inserm, ENS de Lyon, UCBL) facilities: ANIRA IMMOS and especially Laurence Canaple for the metabolic phenotyping, PLATIM and especially Elodie Chatre for the microscopy, for their precious help. We thank our valuable trainees Jules Granet and Emma Fraillon for their contribution in the setting of the protocols. We also thank Dr. Damien Roussel (CNRS, UMR 5023 - LEHNA, Lyon) for helpful discussions regarding the metabolic activity and Dr. Corinne Leprince (INSERM, U1056 - UDEAR, Toulouse) for helpful discussions regarding the keratinocyte terminal differentiation. Alejandro Gonzalez-Torres is grateful for a postdoctoral fellowship from SECTEI.

