## Supplementary Tables for "MiR-30a-5p alters epidermal terminal differentiation during aging by regulating BNIP3L/NIX-dependent mitophagy"

### Supplementary data

| Gene | Gene ID | Primer | Sequence |
|---|---|---|---|
| *KRT10* | 3858 | Forward | 5′-TCCCCCTGATGTGAGTTGC-3′ |
| | | Reverse | 5'-GAATCTGAATGACCGCCTGG-3′ |
| *IVL* | 3713 | Forward | 5'-GCAGTCATGTGCTTTTCCTCTTG-3′ |
| | | Reverse | 5'-TCCTCCAGTCAATACCCATCAG-3′ |
| *LOR* | 16939 | Forward | 5'-TCATGATGCTACCCGAGGTTTG-3′ |
| | | Reverse | 5'-CAGAACTAGATGCAGCCGGAGA-3′ |
| *TGM1* | 7051 | Forward | 5'-GAGAGCACCACACAGACGAG-3′ |
| | | Reverse | 5'-GGGGTTGTTTCCGATGAGTA-3′ |
| *FLG* | 2312 | Forward | 5'-GCTGGAGTATTTTAGGAGATTCTGG-3′ |
| | | Reverse | 5'-CTAGCCCTGATGTTGATATAGCCA-3′ |
| *KLK7* | 5650 | Forward | 5'-TTGGATCACATCAGATCCTCTCG-3′ |
| | | Reverse | 5'-TAATCTTGTCACCCTGGGCTTC-3′ |
| *AQP9* | 366 | Forward | 5'-GTGAGGACCACAACAGGTAGG-3′ |
| | | Reverse | 5'-GCCACATCCAAGGACAATCAAG-3′ |
| *CDSN* | 1041 | Forward | 5'-TCTCCTCCTGCCAGGGAC-3′ |
| | | Reverse | 5'-CGTTAGGGGAGGTGATACGC-3′ |
| *BNIP3L* | 665 | Forward | 5'-TTGGATGCACAACATGAATCAGG-3′ |
| | | Reverse | 5'-TCTTCTGACTGAGAGCTATGGTC-3′ |
| *TBP* | 6908 | Forward | 5'-TCAAACCCAGAATTGTTCTCCTTAT-3' |
| | | Reverse | 5'-CCTGAATCCCTTTAGAATAGGGTAGA-3' |
| *RPL13A* | 6218 | Forward | 5'-CTCAAGGTCGTGCGTCTGAA-3' |
| | | Reverse | 5'-TGGCTGTCACTGCCTGGTACT-3' |
| *HBB* | 3043 | Forward | 5'-CATCAAGCGTCCCATAGACTC-3′ |
| | | Reverse | 5'-ACGTGGATGAAGTTGGTGGT-3′ |
| *SERPINA1* | 5265 | Forward | 5'-AAGGTGAGATCACCCTGACG-3′ |
| | | Reverse | 5'-GTCAGTGAATCACGGGCATC-3′ |
| *MTND1* | 4535 | Forward | 5'-CAGAGACCAACCGAACCCC-3′ |
| | | Reverse | 5'-GAAGAATAGGGCGAAGGGGC-3′ |
| *MTTL1* | 4567 | Forward | 5'-CACCCAAGAACAGGGTTTGT-3′ |
| | | Reverse | 5'-TGGCCATGGGTATGTTGTTAA-3′ |
| **Mutagenesis** | | **Primer** | **Sequence** |
| *BNIP3L* | **Mut1** | Forward | 5'-TGAATTAATGTACAGTCTTCCCAAGGTGATTCC-3′ |
| | | Reverse | 5'-CTGTACATTAATTCAGTGAGAGATCAGAAGGC-3′ |
| *BNIP3L* | **Mut2** | Forward | 5'-TTTACACCAATTTGGGGACAAAAAGGCAGGC-3′ |
| | | Reverse | 5'-CCAAATTGGTGTAAAGCTTTTTTAGC-3′ |
| *BNIP3L* | **Mut3** | Forward | 5'-TATATACAAATACATGTATAACTTGTAGCTATA-3′ |
| | | Reverse | 5'-ATGTATTTGTATATAAAGCCTGCCTTTTTGT-3′ |

**Supplementary Table S1.** List of primers pairs used for qPCR analysis and mutagenesis by reverse PCR.

| Target species | Host species | Dilution | Reference | Provider |
|---|---|---|---|---|
| human BNIP3L | rabbit | 1:200 | HPA015652 | Sigma-Aldrich |
| human KRT14 | mouse | 1:400 | ab7800 | Abcam |
| rabbit IgG | goat | 1:1000 | A-11035 | Thermo Fisher Scientific |
| mouse IgG3 | goat | 1:1000 | 115-605-209 | Jackson ImmunoResearch Ltd |

**Supplementary Table S2.** List of antibodies used for immunofluorescence.
